# Distinguishing discrete and continuous behavioral variability using warped autoregressive HMMs

**DOI:** 10.1101/2022.06.10.495690

**Authors:** Julia C. Costacurta, Lea Duncker, Blue Sheffer, Winthrop Gillis, Caleb Weinreb, Jeffrey E. Markowitz, Sandeep R. Datta, Alex H. Williams, Scott W. Linderman

## Abstract

A core goal in systems neuroscience and neuroethology is to understand how neural circuits generate naturalistic behavior. One foundational idea is that complex naturalistic behavior may be composed of sequences of stereotyped behavioral syllables, which combine to generate rich sequences of actions. To investigate this, a common approach is to use autoregressive hidden Markov models (ARHMMs) to segment video into discrete behavioral syllables. While these approaches have been successful in extracting syllables that are interpretable, they fail to account for other forms of behavioral variability, such as differences in speed, which may be better described as continuous in nature. To overcome these limitations, we introduce a class of warped ARHMMs (WARHMM). As is the case in the ARHMM, behavior is modeled as a mixture of autoregressive dynamics. However, the dynamics under each discrete latent state (i.e. each behavioral syllable) are additionally modulated by a continuous latent “warping variable.” We present two versions of warped ARHMM in which the warping variable affects the dynamics of each syllable either linearly or nonlinearly. Using depth-camera recordings of freely moving mice, we demonstrate that the failure of ARHMMs to account for continuous behavioral variability results in duplicate cluster assignments. WARHMM achieves similar performance to the standard ARHMM while using fewer behavioral syllables. Further analysis of behavioral measurements in mice demonstrates that WARHMM identifies structure relating to response vigor.

## 1 Introduction

A fundamental question in systems neuroscience is how neural activity generates complex behavior [1–3]. Specifically, a key goal is to understand how changes in neural activity determine which actions are selected or executed on a moment-by-moment basis. To make progress towards this goal, it is essential to study ethologically relevant, naturalistic behavior. A common way of studying naturalistic behavior is to observe animals as they freely explore an environment [4–6]. Such unconstrained, spontaneous behavior offers a rich setting for studying behavioral variability. However, the hours of raw video data required to capture naturalistic behavior in detail are high-dimensional and difficult to use for follow-up analyses. Thus, a key goal in behavioral neuroscience research is to develop data-analysis strategies that can extract interpretable lower-dimensional summaries of behavior [4–10]. Ultimately, such descriptions will facilitate relating complex naturalistic behaviors to the underlying neural activity patterns that generate them [3, 11]. This will provide insight into how and why behavior may differ across environmental contexts [2, 3, 11, 12], under pharmacological manipulations [6] or across health and disease [13, 14].

One hypothesis is that the brain generates complex behaviors by concatenating a series of simpler, stereotyped actions [15, 16, 4, 5, 11]. Just like syllables form the building blocks of spoken language, behavioral syllables may be composed to perform complex sequences of behavior. A large focus of previous work has been to discover such behavioral syllables in an unsupervised manner, thereby obtaining a low-dimensional description of high-dimensional behavior [4, 5, 12, 16–19].

Autoregressive Hidden Markov Models (ARHMMs) are well-suited to this task [5, 6]. For example, Wiltschko et al. [5] used depth video to capture the posture of freely moving mice (Fig. 1A). In this work, authors projected the video frames onto the top *D* principal components (PCs) to obtain a *D*-dimensional time series of behavior. The authors then used an ARHMM to segment the behavioral time series into discrete syllables (Fig. 1B). Each discrete syllable corresponds to vector autoregressive dynamics in PC space, which can be conceptualized as a vector field (Fig. 1C). Each instance of a syllable corresponds to a short, stereotyped trajectory following this vector field (red-to-blue trajectories in Fig. 1C). Empirically, these often correspond to stereotyped patterns of movement like rearing, darting, or grooming (Fig. 1D).

**Figure 1:**
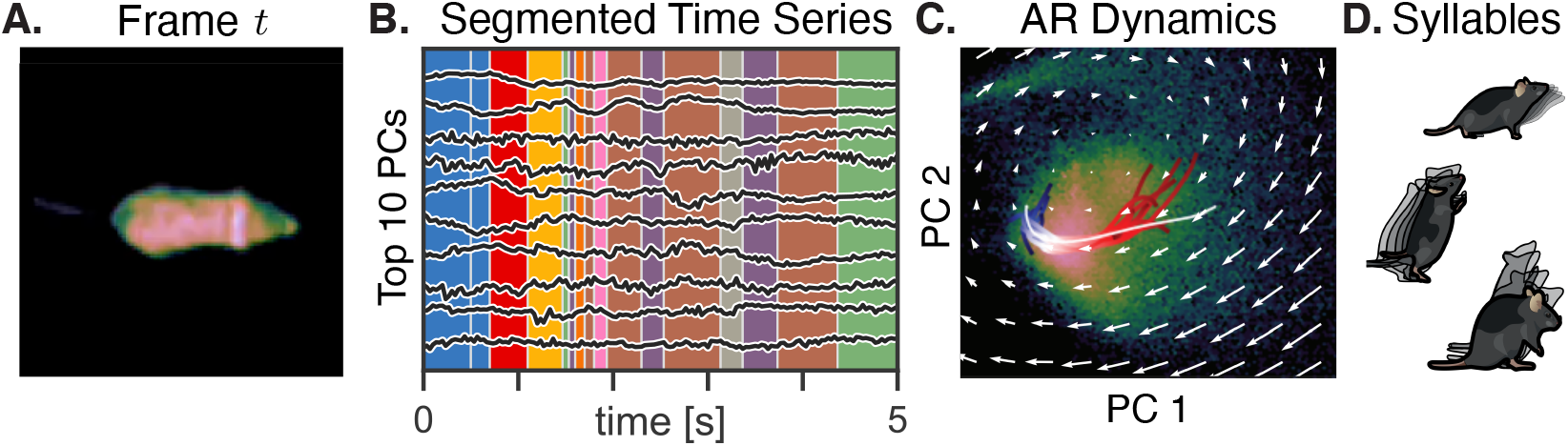
**A**. Single frame of depth video of a freely behaving mouse. **B**. The frames are projected onto their top ten principal components (black lines) and then segmented into discrete syllables (colors) with an autoregressive HMM (ARHMM). **C**. Each syllable is defined by autoregressive (AR) dynamics, which can be visualized as a flow field in PCA space. Here, the scatter plot shows the distribution of frames projected onto the top two PCs, and the red-to-blue trajectories correspond to multiple instances of one discrete state. **D**. Empirically, syllables correspond to interpretable behaviors like investigating, rearing, and falling from a rear, as illustrated here.

However, current modeling approaches have focused on clustering *discrete* behavioral syllables while failing to account for other, *continuous* forms of behavioral variability. For example, the same type of behavior (e.g. a dart) could be performed more or less vigorously. While these two actions might be identified by a human observer as the same behavior under different speeds (fast vs. slow darts), current models lack the ability to allow for such structured continuous variability within states, and thus might assign these actions to entirely separate behavioral states. As a result, current approaches often *over-segment* video data, by allocating distinct clusters to the same movement type. The data might be better described by a model which incorporates a continuous spectrum of structured variability within a specific behavior.

We extend the ARHMM by incorporating a latent *warping variable*. With this Warped Autoregressive Hidden Markov Model (WARHMM), we are able to capture continuous variability within a discrete syllable. Thus, we are able to disentangle variability due to (discrete) movement type from other forms of (continuous) variability, such as movement speed. We consider two types of warping: *time-warping*, which captures changes in the speed of evolution of the autoregressive dynamics, and *Gaussian process warping*, which allows for nonlinear changes in dynamics. We develop an efficient inference algorithm for the WARHMM and show that it can correctly identify the underlying states and parameters in simulated data. Then, using behavioral measurements of freely behaving mice, we demonstrate that accounting for continuous sources of behavioral variability can resolve issues of over-segmentation commonly observed using previous approaches. Furthermore, using behavioral data of mice treated with either saline solution or amphetamine, we demonstrate that the WARHMM identifies differences in the distribution of latent warping variables across both groups of mice. This result reflects potential modulations of movement vigor and speed distributions due to the pharmacological intervention.

## 2 Background

We first review related work and then present a brief description of two classes of Hidden Markov Models that have been used for time series segmentation and behavioral modeling. The key features of each model form the basis for our warped extension to the ARHMM model, which represents the main contribution of our work.

### Related work in unsupervised behavioral segmentation

We build upon a rich body of work in the unsupervised behavioral segmentation space. Our model is inspired by the ARHMM approach to unsupervised behavioral segmentation (MoSeq) proposed by Wiltschko et al. [5], which is described in the introduction. Berman et al. [4] addressed a similar task in fruit fly video using MotionMapper, which identifies discrete behavioral states as peaks in a non-linear, two-dimensional embedding of postural spectrograms. Hsu and Yttri [17] have proposed B-SOiD, which uses a random forest to classify non-linear postural feature embedding clusters into multiple behavioral classes. Harris et al. [18] fit an autoregressive linear model to time-windowed postural features using low-rank tensor decomposition, and interpret clusters in the fit model parameters as discrete behavioral states. Luxem et al. [19] identify behavioral states by clustering the latent vectors produced by training a variational autoencoder on input from markerless pose estimation [9]. For a more thorough treatment of work in unsupervised behavioral quantification, see McCullough and Goodhill [20] for a recent review. The majority of these related approaches have focused on segmenting behavioral video based on movement type or on tracking animal pose. The model we introduce in section 3 extends on this work by explicitly taking other forms of behavioral variability, such as movement speed, into account.

### Autoregressive Hidden Markov Models

An ARHMM (Fig. 2A, top) consists of a discrete latent state variable *z*_*t*_ ∈ {1, 2, …, *K*} and observations **x**_*t*_ ∈ ℝ^*D*^. Transitions between the discrete state values over time are governed by a transition matrix **P**, where the entry **P**_*k,k*_′ indicates the probability of advancing from state *z*_*t*_ = *k* to state *z*_*t*+1_ = *k*^′^. Given the discrete state value, the observations are modeled to evolve according to linear dynamics, where a dynamics matrix 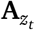 and bias term 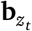 determine the mapping from **x**_*t*_ to **x**_*t*+1_ in the presence of Gaussian noise with covariance matrix 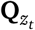. Thus, the model represents an extension of classic Gaussian mixture models to time series with autoregressive dynamics. The ARHMM model can be summarized as

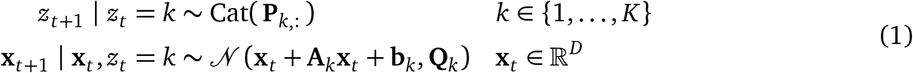

**Figure 2:**
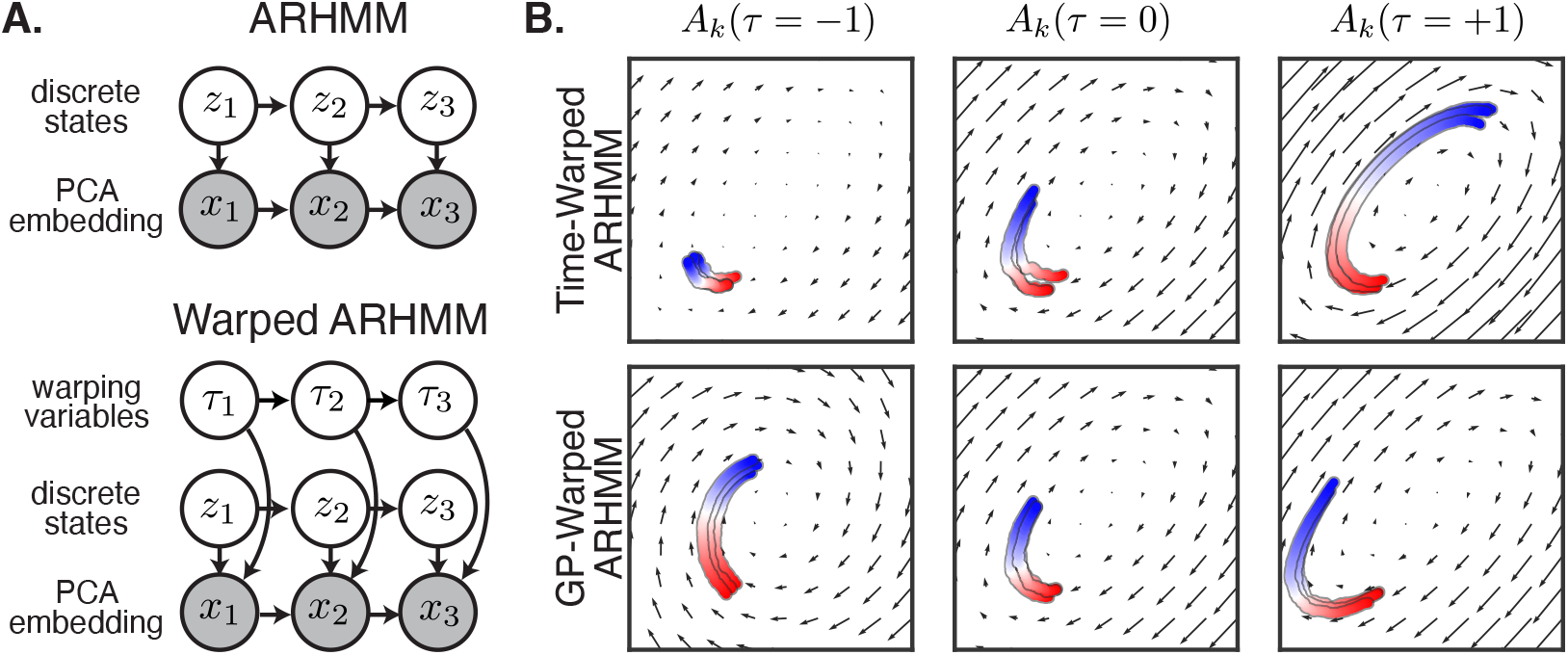
**A**. Probabilistic graphical models for the ARHMM and the Warped ARHMM. The warping variables and discrete states (i.e. syllables) together determine the AR dynamics. **B**. We consider two types of warping: time-warping, where dynamics are sped up or slowed down by the warping variable, and GP warping, where the dynamics vary smoothly but nonlinearly with the warping variable.

In the context of behavioral modeling [5], the discrete latent state reflects the behavioral syllable (such as a dart, leftward turn, rear, etc.) and determines which dynamics are used to describe the temporal evolution of the observed posture **x**_*t*_.

### Factorial Hidden Markov Models

In Factorial Hidden Markov Models (FHMMs) [21], multiple discrete hidden state variables influence the distribution of observed states. The FHMM model can be summarized as

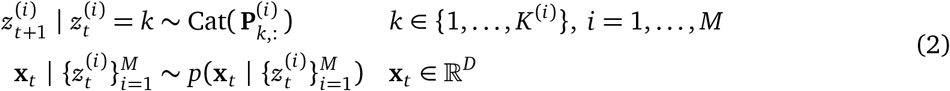

FHMMs are useful in cases when aspects of data can be distributed across multiple states. For example, to fit data that varies according to *M* binary variables, a standard HMM would need 2^*M*^ discrete states, while a FHMM would only need *M* binary states [21].

In the context of behavior, adding additional latent variables is an appealing way to provide structured variability within discrete syllables. Assuming this variability is included in a reasonable way, an FHMM has the potential to describe a dataset with fewer discrete latent states than an HMM.

## 3 Warped Autoregressive Hidden Markov Models

ARHMMs have been successful in clustering video measurements of behavior into discrete sets of behavioral syllables. However, in practice they are prone to *over-segmentation*, where behaviors that appear similar to a human expert are split into distinct clusters. We hypothesize that oversegmentation arises because the ARHMM conflates discrete sources of behavioral variability — the expression of distinct behavioral syllables — with continuous sources of variability that cannot be captured by linear autoregressive dynamics with Gaussian noise.

To address this limitation, we develop Warped Autoregressive Hidden Markov Models (WARHMMs). WARHMMs extend the ARHMM with an additional latent variable, like in a Factorial HMM. In addition to the discrete state variable *z*_*t*_, WARHMM includes a latent warping state *τ*_*t*_ ∈ [−1, 1] at each time step (Fig. 2A, bottom). The warping state *τ*_*t*_ modulates the dynamics associated with each state *z*_*t*_. While *z*_*t*_ can model rapid switches in dynamics, *τ*_*t*_ can account for additional variability in the dynamics associated with a given latent state.

The general form of this model class can be summarized as follows, where **A**_*k*_(*τ*_*t*_), **b**_*k*_(*τ*_*t*_), and **Q**_*k*_(*τ*_*t*_) are functions of *τ*_*t*_ :

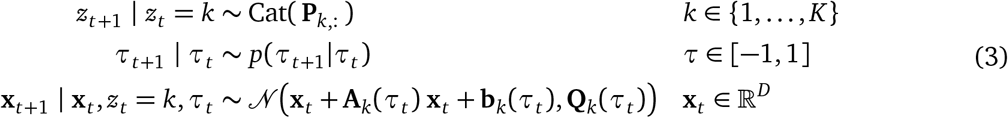

Below, we consider two possibilities for how exactly the warping variable modulates the dynamics.

### 3.1 Time-Warped ARHMM: linear modulation of autoregressive dynamics

Our first specific instance of WARHMM has a direct motivation in terms of time-warping, and we thus refer to it as a time-warped ARHMM (T-WARHMM). Specifically, we aim to capture continuous changes in how quickly trajectories move through observation space. If we consider the change in current state Δ**x**_*t*_ = **x**_*t*+1_ − **x**_*t*_ due to the dynamics within a given state *z*_*t*_ = *k* and let *τ* be a *log step-size parameter*, we can write

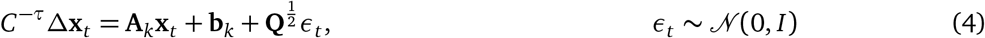

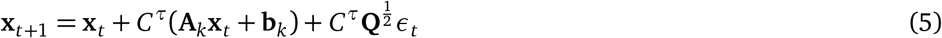

When *τ* = 0 the step size *C*^*τ*^ is always one, so equation (5) is equivalent to the classic ARHMM. For nonzero *τ*, however, the warping variable modulates how far a single update can move the state, akin to a time-constant in ordinary differential equations. The constant *C* determines the maximum multiplicative factor by which dynamics can be scaled; in our experiments we set *C* = 2. In terms of the functional mapping, T-WARHMM corresponds to the following mapping between the warping state *τ*_*t*_ and the parameters determining the autoregressive dynamics of the observed states **x**_*t*_ :

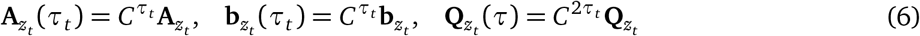

Fig. 2B (top) shows how an example syllable’s dynamics are sped up or slowed down by changing *τ*. As *τ* increases, the trajectories traverse a greater distance in the same number of time steps. In the context of behavioral modeling, the time-warping variable may capture modulations in movement speed. As we will see in section 5, the inferred warping variables correlate with other intuitive notions of vigor, like centroid velocity in darting syllables, while also offering a quantification of vigor in stationary syllables such as grooming.

### 3.2 Gaussian Process-WARHMM: Nonlinear modulation of autoregressive dynamics

The model formulation for T-WARHMM is interpetable and intuitive, but also makes strong parametric assumptions about how continuous variability affects the dynamics of behavior. As a point of comparison, we consider a more flexible model, in which the influence of *τ*_*t*_ via 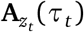 and 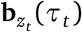 is specified nonparametrically in terms of Gaussian processes. This Gaussian Process WARHMM (GP-WARHMM) moves beyond using *τ*_*t*_ to describe speed or vigor, as *τ*_*t*_ could have effects beyond modulation of time-constants of the dynamics. Here *τ*_*t*_ can modulate entries of the dynamics matrix according to a smooth nonparametric function. In particular, the dynamics are modeled as follows:

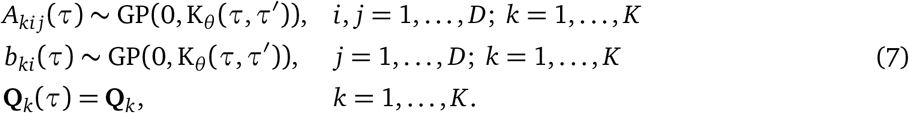

where *A*_*ki j*_(*τ*) denotes the (*i, j*)-th entry of the matrix **A**_*k*_(*τ*) and *b*_*ki*_(*τ*) is the *i*-th entry of the vector **b**_*k*_(*τ*). Thus, each coordinate varies smoothly and independently as a function of *τ, a prioir*. In our experiments, we choose a squared exponential kernel with kernel hyperparameters *θ* = (*ρ, σ*).

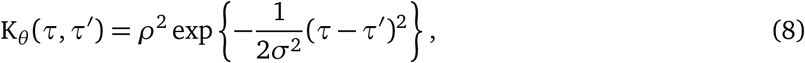

Fig. 2B (bottom) shows how an example syllable’s dynamics are modulated by changing *τ* in a GP-WARHMM. The central dynamics (*A*_*k*_(*τ* = 0)) are the same as in the T-WARHMM above, but now *τ* can have nonlinear effects on the dynamics that do more than simply slow them down or speed them up. This extra flexibility could be useful for capturing more general types of continuous variability within discrete syllables, but it may also come at the cost of interpretability.

### 3.3 Inference and Learning

Though we have presented *τ*_*t*_ ∈ [−1, 1] as a continuous latent variable, in practice we find it sufficient to discretize *τ*_*t*_ over a fine grid of *J* evenly spaced points, since it is a one-dimensional, bounded random variable. Once discretized, we model the warping variable dynamics as a Markov process with a banded transition matrix to encourage small, local changes over time. Similarly, the GP prior in eq. (7) reduces to a multivariate normal after discretization. Full details are provided in the Appendix.

As in standard ARHMMs, we estimate the states and parameters with the Expectation-Maximization algorithm (EM) [22]. We perform exact inference over the discrete syllables and warping variables using the forward-backward algorithm, which runs in *O*(*T*(*K*^2^ + *J*^2^)) time and uses *O*(*TKJ*) memory. For both the time-warped and GP-warped ARHMMs, the parameter updates have closed-form solutions. For large datasets, we used stochastic EM [23] to speed convergence. For the GP-WARHMM, we manually tuned the kernel hyperparameters, but they can in principle be learned by type-II maximum likelihood learning. Again, complete details are in the Appendix.

## 4 Synthetic data validation

We begin by generating two-dimensional synthetic data **x**_*t*_ ∈ ℝ^2^ with *K* = 2 discrete states. The autoregressive dynamics under each discrete state are chosen to be a clockwise and counterclockwise rotation. We modulated the dynamics with a time-warped ARHMM, setting **A**_*k*_(*τ*) = 2^*τ*^**A**_*k*_, as illustrated in Fig. 3A. For this simple experiment, we limited the true time-warping variables to *J* = 5 values of *τ* evenly spaced on [−1, 1]. A snippet of simulated data and the corresponding time constants and states is shown in Fig. 3B. We fit both a GP-WARHMM and T-WARHMM with *K* = 2 states and *J* = 5 warping variables to data simulated from this model.

**Figure 3:**
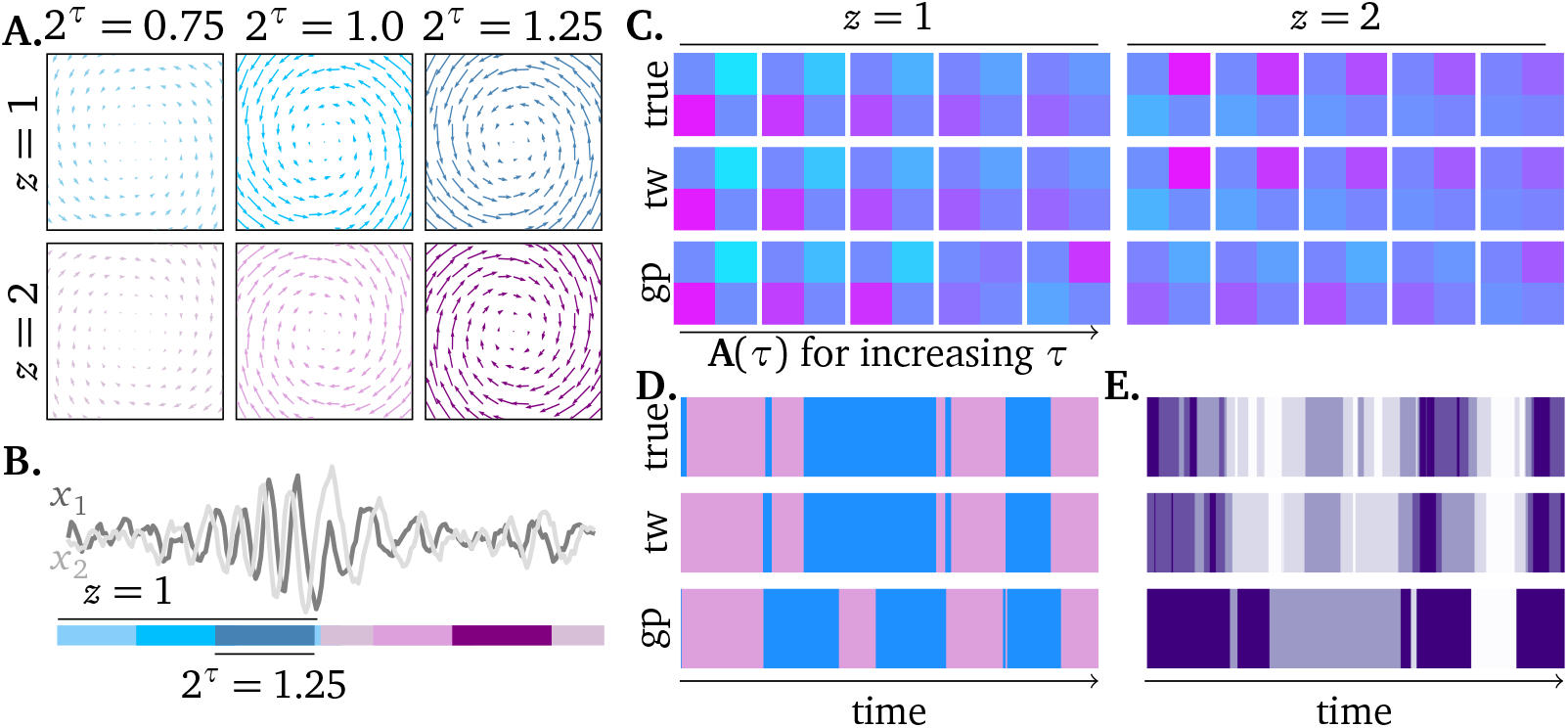
**A**. Illustration of the dynamics of the simulated example. Two discrete latent states represent a clockwise (*z* = 2, bottom) and counterclockwise (*z* = 1, top) rotation, each shown as vector fields. The speed of the rotation is modulated via different values of *τ*, where **A**_*z*_(*τ*) = 2^*τ*^**A**_*z*_. **B**. Illustration of the state trajectory as the generative dynamics are modulated by both state switches in *z* (discrete variability) and changes in rotation angle via *τ* (continuous variability). **C**. The generative value for **A**_*k*_(*τ*) = 2^*τ*^**A**_*k*_ (top row) together with the learned matrices for T-WARHMM (tw, middle) and GP-WARHMM (gp, bottom). T-WARHMM is able to learn the variations in the autoregressive dynamics. GP-TWARHMM learns the overall rotation structure of this examble, but the increased flexibility of the GP model makes it harder for it to disentangle discrete and continuous contributions to changes in **A**_*k*_(*τ*). **D**.-**E**. The true and inferred latent paths (posterior mode) for *z*_*t*_ (**D**) and *τ*_*t*_ (**E**) under each model.

The results from this experiment are summarized in Fig. 3C-E. Panel C shows the true and learned values of entries of the autoregressive dynamics **A**_*z*_(*τ*) as both *z* and *τ* vary. We see that both the T-WARHMM and GP-WARHMM learn dynamics that reflect the overall rotational structure of the matrix. However, only T-WARHMM is able to recover the clockwise and counterclockwise variation with *z* and correct speed modulation with *τ*.

The increased flexibility of GP-WARHMM, in contrast, allows it to achieve a comparable test log likelihood (Table 1), while not being able to correctly disentangle discrete and continuous contributions to the observed data variability. Fig. 3D and E show the true state changes in *z*_*t*_ and *τ*_*t*_, together with the mode of the inferred posterior distributions under each model. T-WARHMM is able to recover the correct segmentation of the data, while GP-WARHMM’s increased flexibility allows it to fit the data, but in a less interpretable way.

**Table 1:** Comparison of test log likeli-hoods on simulated data across true and fitted models.

| Model | Test LL |
| --- | --- |
| T-WARHMM | 0.0180 |
| GP-WARHMM | -0.0040 |
| True Model | 0.0344 |

## 5 Modeling depth video of freely behaving mice

For the remainder of the paper, we are interested in extracting discrete and continuous structure in behavior from posture data **x**_*t*_ extracted from depth-imaging recordings of freely behaving mice. To do this, we reanalyze data from Wiltschko et al. [5], which represents the original application of the ARHMM to clustering mouse behavior. In the context of this dataset, the ARHMM approach is often also referred to as *MoSeq*, and we thus refer to the dataset as the MoSeq dataset.

In the MoSeq dataset, the observations **x**_*t*_ are taken to be the first 10 principal components of depth camera video data of mice exploring an open field. The dataset consists of 20-minute depth camera recordings of 24 mice. In preprocessing, the videos are cropped and centered around the mouse centroid, and then filtered to remove recording artifacts. Finally, the preprocessed video is projected onto the top principal components to obtain a 10-dimensional time series.

### 5.1 Comparison of model performances on the MoSeq dataset

To provide a direct comparison of performance between the ARHMM, T-WARHMM, and GP-WARHMM, we trained each model using 50 epochs of stochastic EM. The ARHMM is equivalent to setting the number of *τ*-values in T-WARHMM to *J* = 1. T-WARHMM and GP-WARHMM each had *J* = 31 evenly spaced values of *τ* on the interval [−1, 1].

Fig. 4A shows the log-likelihood of each model on held-out test data for a range of *K* values. For a given value of *K*, both T-WARHMM and GP-WARHMM outperform the classic ARHMM in terms of their generalization performance on unseen data. However, we achieve similar test log-likelihoods for an ARHMM with *K* = 40 syllables, a T-WARHMM with *K* = 25 syllables, and a GP-WARHMM with *K* = 15 syllables. This further illustrates that the ARHMM has to account for continuous variability by creating a larger number of discrete states, while the factorial structure of the WARHMMs enables syllables to be merged when the difference in their dynamics can be explained by changes in *τ*. The GP-WARHMM outperforms the T-WARHMM in terms of test log-likelihood, which can be attributed to the fact that the GP-WARHMM is more flexible than the T-WARHMM and can capture additional structure in the data. However, the GP-WARHMM accomplishes this at the cost of interpretability. The T-WARHMM warping constant is physiologically motivated and, as we will show in the later results, has direct correlations to centroid velocity and other notions of vigor. While the GP-WARHMM warping variable is allowed to vary the syllable in whichever way will increase data log-likelihood, the T-WARHMM is restricted to modulating syllables in a way that is biologically interpretable. We believe that this interpretability of T-WARHMM makes it more useful for behavioral analysis purposes. For this reason, we focus on the utility of T-WARHMM for the remainder of this paper.

**Figure 4:**
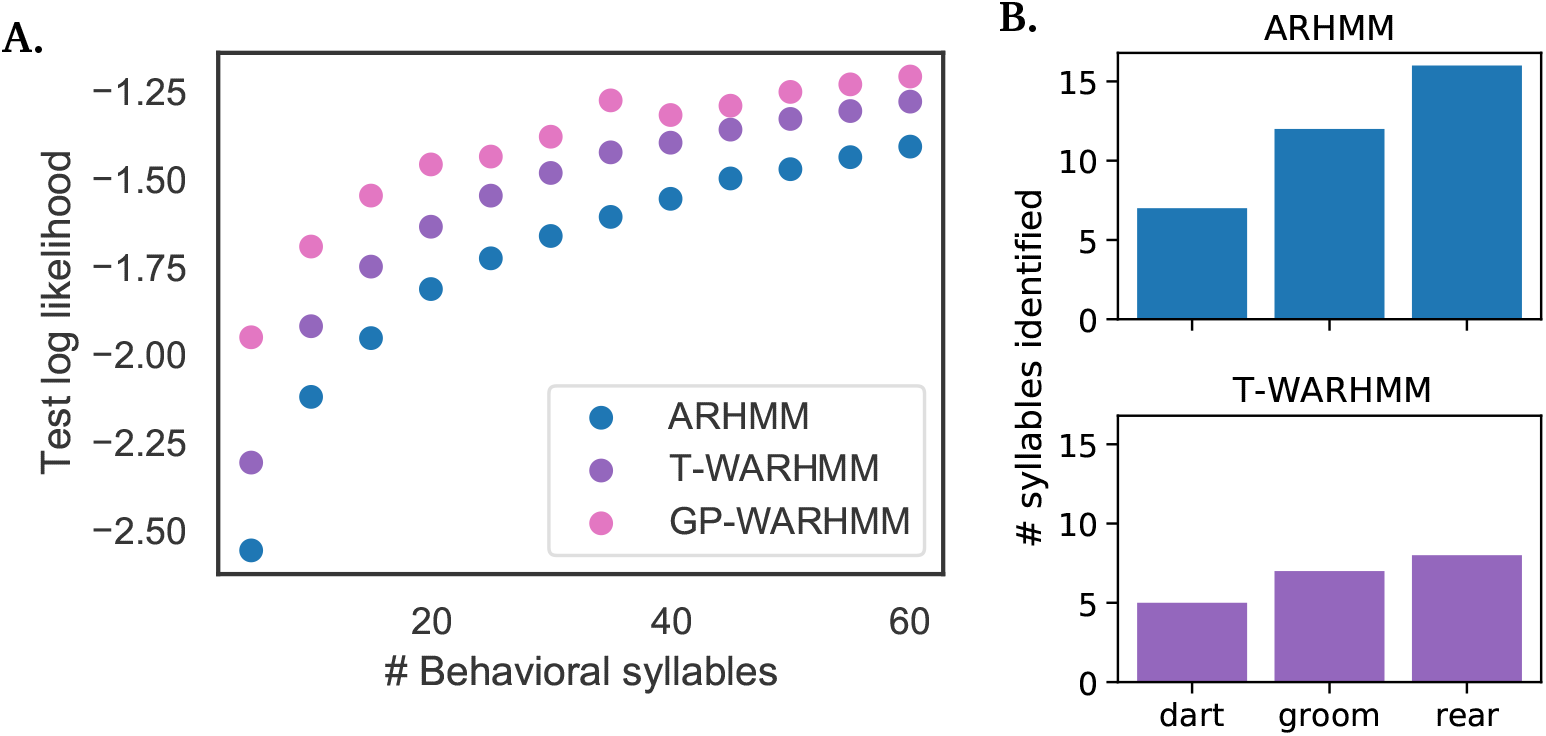
**A**. Comparison of log-likelihood on held-out test data as a function of the number of discrete latent states after training the different model classes on the MoSeq dataset. **B**. For an ARHMM and T-WARHMM with similar test log-likelihoods, total number of syllables that are grouped into the same behavior class by expert neuroethologists. The ARHMM creates new syllables to model qualitatively similar behaviors.

### 5.2 Interpreting the time-warping variable

#### Which syllables are “warped”?

To begin our analysis of the T-WARHMM results, we fit a model with *K* = 20 discrete states and *J* = 31 time constants. As shown in Fig. 4A, this model attains similar test log-likelihood to an ARHMM with *K* = 35 discrete states. We then determine which types of syllables are over-represented as multiple distinct discrete states in the ARHMM by analyzing the syllable labels from each model. After generating sample videos of behavior from both models, we asked expert neuroethologists and MoSeq users to label the syllables produced. We then gathered the labels under three general behaviors: darts, grooms/pauses, and rears. The results are shown in Fig. 4B. Both models have similar numbers of darting states, while the numbers of grooms and rears are reduced in T-WARHMM, with T-WARHMM halving the number of rear states. While T-WARHMM does not completely resolve the oversegmentation issue, it is able to perform as well as the ARHMM with fewer syllables.

#### Connections to centroid velocity

From our formulation of the time warping variable, it is expected that the time warping variable and centroid velocity of the mouse would be highly correlated. In Fig. 5 we plot measured centroid velocity (in pixels/ms) vs. inferred vigor (2^*τ*^) for four representative states. We see a strong relationship between centroid velocity and vigor for the darting state, where we would expect such a relationship to occur. The other states also show this relationship, but less consistently. In these states, the warping variable may be accounting for additional forms of timing-related variability that are not well accounted for by centroid speed. These analyses validate that the time warping parameter *τ* is able to extract speed-related information also contained in centroid velocity for behavioral syllables such as darting motions.

**Figure 5:**
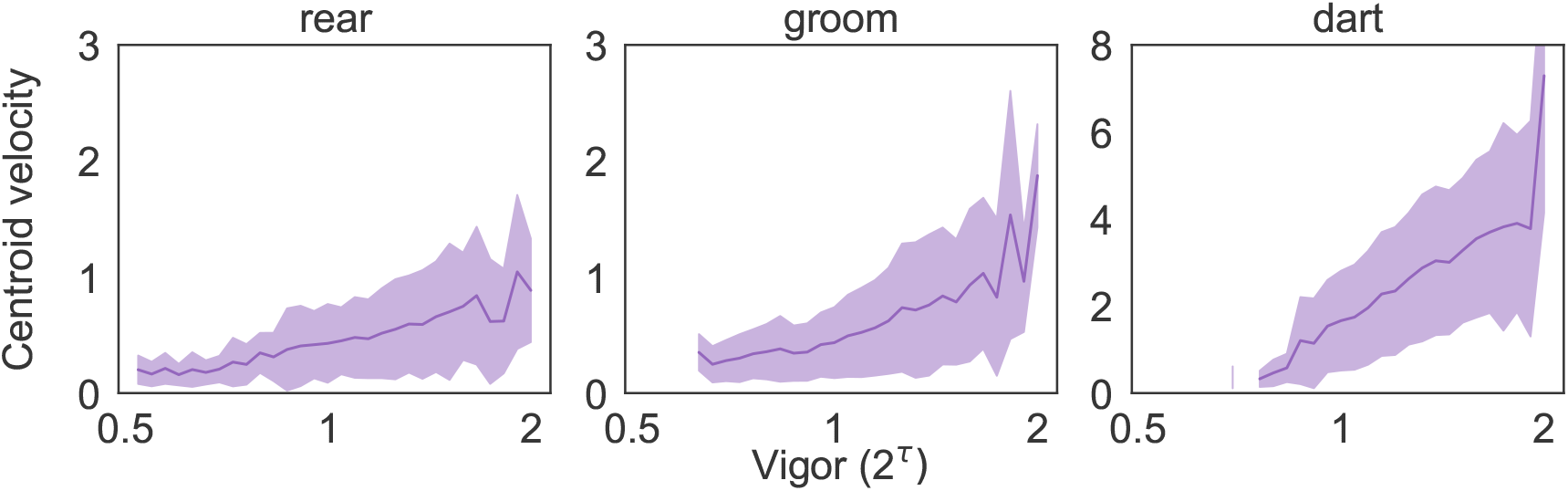
Centroid velocity (pixels/ms) vs. inferred vigor (2^*τ*^) for three representative states of the TW-ARHMM, showing a clear positive correlation between the variables.

### 5.3 Identifying drug-induced changes in behavior

Finally, we analyze video recordings of two groups of mice treated with either saline solution or amphetamine. As shown in Fig. 6A, the distribution of centroid speed varies slightly between the groups, with the amphetamine-treated mice performing actions at higher speeds more frequently than the saline mice. We fit a T-WARHMM with *J* = 31 time constants and *K* = 20 syllables to both sets of data. Fig. 6B shows the the inferred vigor (2^*τ*^) distributions from two representative syllables, demonstrating that there is a clear rightward shift between amphetamine and saline treated mice. This indicates that amphetamine mice use faster time warping variables more frequently than saline mice. The difference between means of the *τ* index distributions are shown in Fig. 6C. Our model shows a significant difference (*p* < 0.05, independent t-test) between the means of the *τ* distributions for all except one of the inferred syllables. Thus, T-WARHMM is able to detect and dynamically track drug-induced differences in behavior across both groups of mice.

**Figure 6:**
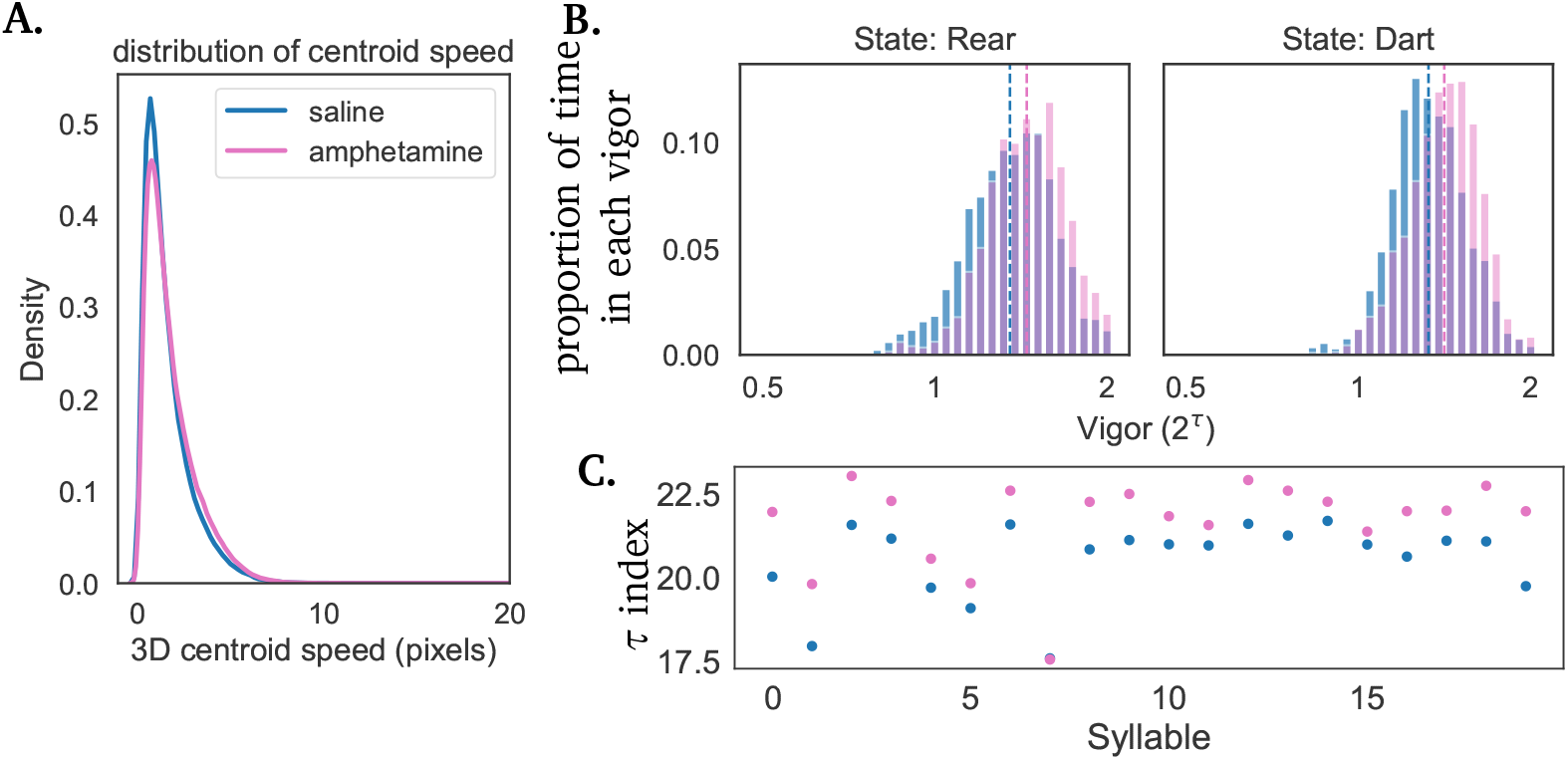
**A**. The distribution of centroid speed across mice treated with saline (blue) or amphetamines (pink). **B**. Histograms showing the vigor distributions across the two classes of mice treated with saline (blue) or amphetamines (pink) for two example syllables (discrete states). The dashed lines indicate the mean of distributions, showing a clear increase in mean vigor in the amphetamine treated group. **C**. Mean *τ* index for each of the 20 discrete states in the model for both the saline (blue) and amphetamine (pink) treated mice. The inferred *τ* indices in the amphetamine treated group are larger (higher vigor, faster movement speed) for all but one syllable.

## 6 Conclusion

We have introduced an approach for unsupervised behavioral modeling which allows for the identification of continuous variability within discrete behavioral syllables. Our work builds on a prior state-of-the-art technique, which uses a discrete hidden state in combination with autoregressive linear dynamics to cluster behavior. In our proposed model extension (WARHMM), underlying the behavioral measurement at each time point are both a discrete state variable and a warping variable. Both of these latent variables modulate the dynamics of the syllable, either through discrete switches (via the discrete state variable) or through more continuous forms of modulation (via the warping variable). We found that WARHMM achieved similar performance in terms of log-likelihood on held out test data while utilizing fewer discrete latent states than the ARHMM.

Our analyses on mouse behavioral data focused on an interpretable subclass of warped ARHMM, where the warping variable interacts linearly with the autoregressive dynamics of the observed data (the T-WARHMM). Notably, the warping variable in this simple model can be viewed as implementing a local rescaling of time. Similar time warping models have been fruitfully applied to trial-structured neural data [24, 25], and our work illustrates how these ideas can be extended to naturalistic time series data that lack repeated trial-structure. Our work on mouse behavioral data illustrates that continuous variability in behavior may be induced through pharamacological interventions and can be extracted directly from video recordings.

### Limitations

For simplicity, we discretized *τ* on a fine grid instead of explicitly performing inference over the continuous variable. This was done in favor of computational efficiency and ease of inference. While we believe the fine grid we’ve used for *τ* offers a good approximation of a continuous function, future work could capture continuous variability by including a truly continuous posterior, e.g. via a Gaussian Process [24]. Furthermore, our model assumes that the behavioral PCs evolve according to linear dynamics, and though the GP-WARHMM allows these dynamics to be modulated in nonlinear ways, the basic assumption could be relaxed to more general nonlinear dynamics in future work.

### Future work

This work establishes multiple directions for future research. The first involves further extending the ARHMM model by adding in additional forms of structured, interpretable variability. What aspects of behavior, in addition to speed, may vary within syllables? In the context of mouse behavior, variables such as turning radius or depth-camera pixel height may provide additional continuous axes along which behaviors can be modulated. Another future direction lies in using T-WARHMM to analyze neural correlates of behavior. Similar behavioral segmentation approaches have been used to simultaneously analyze the evolution of behavioral and neural recordings [26, 8, 27, 28]. A future research area of particular interest lies in understanding how the dorsolateral striatum encodes action selection and vigor [29, 13, 30–32]. Neural activity in this midbrain nucleus is known to be disrupted in motor diseases like Parkinson’s and Huntington’s [33, 34]. The T-WARHMM approach allows us to simultaneously extract descriptions of movement type (how frequently actions are selected) and movement speed (how vigorously actions are performed) from behavioral measurements. We expect that the ability to disentangle the contributions of these two factors could provide an important basis for better understanding how variations in neural activity determine changes in action selection and vigor. Finally, the models we have proposed here could be applied more widely to analyze other types of behavioral data.

## Acknowledgements

We would like to thank Andy Warrington, Laura Driscoll, Libby Zhang, Jimmy Smith, Michael Salerno, Mohammed Osman, Sherry Lin, and Akshay Jaggi for their feedback and suggestions.

This work is supported by the Simons Collaboration on the Global Brain and NIH BRAIN Initiative Grant 1U19NS113201-01. JC is funded by the NSF GRFP and Stanford Graduate Fellowship.

## Author Contributions

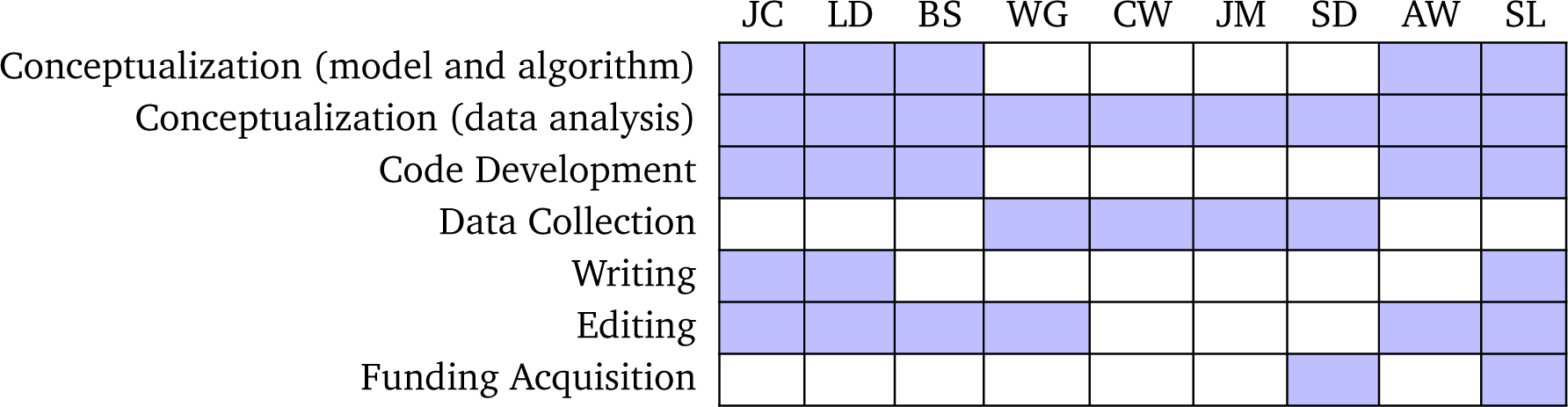

## A Inference and Learning

We fit warped ARHMMs to behavioral measurements using the Expectation-Maximization (EM) algorithm. Here, we provide additional details on performing posterior inference over the latent variables *z*_1:*T*_ and *τ*_1:*T*_ (E-Step), and include details on closed-form parameter updates (M-Step).

### A.1 Inference

Our inference approach is the same for both model classes of WARHMM presented in the main text. Inference is performed using forward-backward message passing, with a slight twist to speed inference over a large number of (*z, τ*) hidden state pairs. In particular, during message passing the *K* discrete states and *J* warping variables are represented as *K* · *J* “paired” states, so inference via message passing becomes very slow due to multiplication with a *KJ* × *KJ* transition matrix. To ease this bottleneck we enforced Kronecker structure on the *KJ* × *KJ* transition matrix:

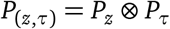

Then, if we let *α*_*t*_, *β*_*t*_ ∈ ℝ^*K*×*J*^ be the forward and backward messages defined on the grid of (*z, τ*) values, the recursive calculations become:

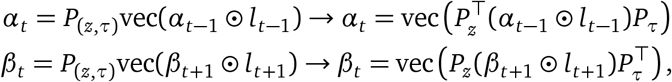

where ⊙ denotes elementwise multiplication, *l*_*t*_ ∈ ℝ^*K*×*J*^ denotes the likelihoods over the grid of (*z, τ*) values at time *t*, and vec(·) denotes the vectorization operation which concatenates columns of a matrix into a single column vector. This results in a complexity decrease of *O*(*K*^2^*J*^2^) to *O*(*K*^2^ + *J*^2^) in the matrix-vector multiplication. The posterior marginal distributions are proportional to the elementwise product, *q*(*z*_*t*_, *τ*_*t*_) ∝ *α*_*t*_ · *l*_*t*_ · *β*_*t*_.

### A.2 Learning

In this section, we provide additional details on the parameter updates under each model class.

#### A.2.1 T-WARHMM

The closed form updates for the model parameters under the T-WARHMM model class are given by

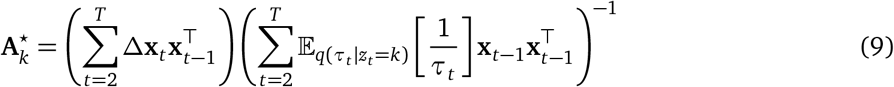

and

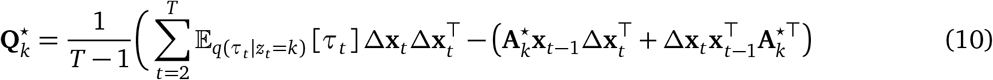

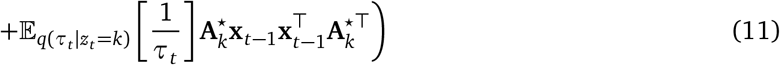

where Δ**x**_*t*_ = **x**_*t*_ − **x**_*t*−1_.

#### A.2.2 Gaussian Process WARHMM

We place independent Gaussian priors over the transition dynamics

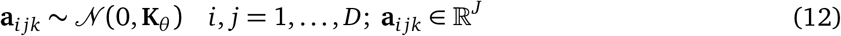

The prior covariance is constructed via an exponetiated quadratic covariance function such that

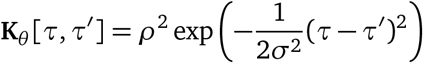

is evaluated on the grid of *τ* values and *θ* = {*ρ, σ*}. The full set of autoregressive dynamics for the different (*z*_*t*_, *τ*_*t*_) pairs, **A**, is a *D* × *D* × *K* × *J* tensor. We will work with different reshapings of this tensor to obtain to derive a closed form update.

Let 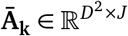 denote the reshaping of the transition dynamics. Here, each column corresponds to the vectorization of a *D* × *D* slice of the tensor, and each column corresponds to a different value on the *τ* grid. The prior over **Ā**_**k**_ can be expressed as a Matrix Normal distribution with identity row covariance *I* and column covariance **K**_*θ*_ :

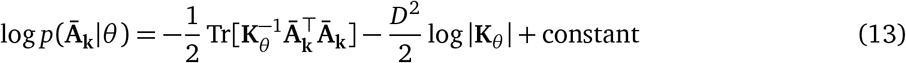

To find the update for **Ā**_**k**_, we need to maximize

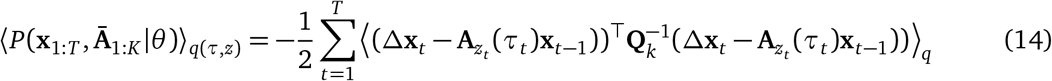

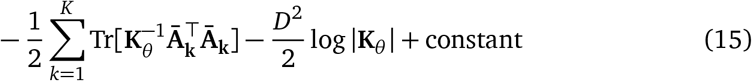

We can rewrite 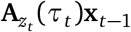 as

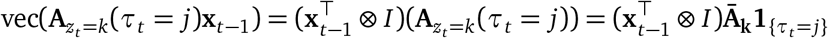

where 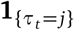 is a length-J vector with binary entries, which selects the column from **Ā**_**k**_ for which *τ*_*t*_ = *j*.

Letting *q*(*z*_*t*_ = *k, τ*_*t*_ = *j*) = *ω*_*kjt*_, differentiating with respect to **Ā**_**k**_ and setting to zero, we obtain:

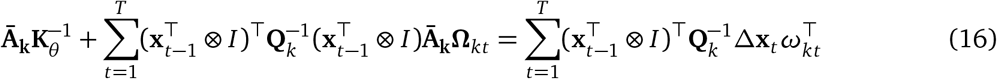

where **Ω**_*kt*_ = diag(*ω*_*k*1*t*_, …, *ω*_*kJt*_) = diag(*ω*_*kt*_). We can show that

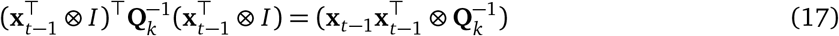

and

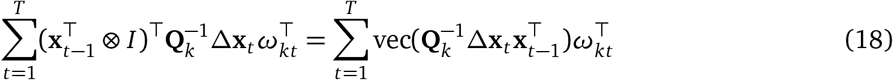

Substituting these expressions into Equation (16), we obtain

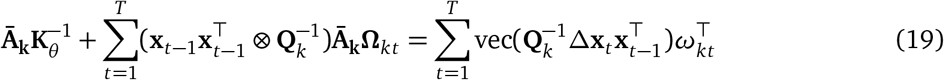

To solve this equation for **Ā**_**k**_ in closed form, we can rewrite the above in terms of the the length *D*^2^*J* vector **ā**_*k*_ = vec(**Ā**_**k**_). We obtain the update equation

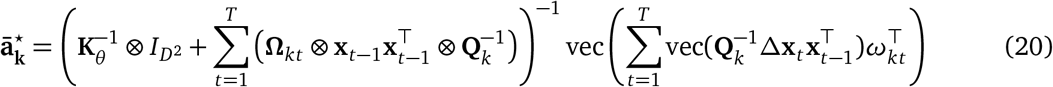

The update above can be expressed more efficiently in terms of expected sufficient statistics. Letting 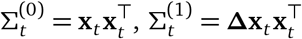 and 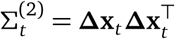, we obtain the expected sufficient statistics

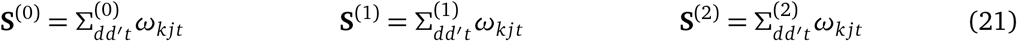

where we have used tensor index notation to indicate a summation over the time index. **S**^(0),(1),(2)^ are *D* × *D* × *K* × *J* tensors. Precomputing and storing these statistics after each E-step allows for efficient M-Step updates.

The update for **Q**_*k*_ takes the closed form

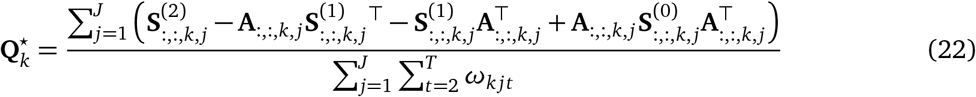

## B Computing details

### B.1 Code

The code required to reproduce our main results is published at https://github.com/lindermanlab/warhmm.

### B.2 Training details

All of the models shown in the paper results were trained with 50 epochs of either EM (simulated data) or stochastic EM (MoSeq dataset). The data was split 80/20 into train and test datasets.

#### Hyperparameters

We set specific hyperparameters as follows:

1. *κ* and *α* are as in [5] and enforce a prior on discrete state transitions, so that discrete states are more likely to remain the same for multiple time steps before switching. In our code, based on a previous sweep over *α* and *κ* we found that *α* = 5 and *κ* = 10000 produced discrete states with durations long enough to be interpretable as behavioral syllables.
2. *τ*_stay_ is the probability that *τ*_*t*_ = *τ*_*t*+1_, or the diagonal entry of the *τ* transition matrix. To approximate *τ* as continuous, we enforced a banded structure on the transition matrix so *τ* could only transition to adjacent values between time steps, and the off-diagonal values were (1 − *τ*_stay_)*/*2. In our training we tried *τ*_stay_ = 0.7 and *τ*_stay_ = 0.9. We found that *τ*_stay_ = 0.7 allowed for more variation of *τ* within discrete states, matching our desire that the warping variable could change its effect while a discrete state is carried out. Thus, *τ*_stay_ = 0.7 was used for the paper results.
3. *σ* and *ρ* are the two parameters of the GP-WARHMM’s RBF kernel. We tried three values of the lengthscale *σ* in our experiments: 0.5, 1, 2. We found that there was little difference in test log-likeihood between these values of *σ*, so the experiments in the paper use *σ* = 1 to encourage smoothness in the learned GP functions without being too restrictive. Due to compute time we did not perform a similar sweep over the output scale *ρ*, and took *ρ* = 1 in the paper experiments.
4. *C* is the log step-size parameter for the T-WARHMM warping variables. In the paper experiments we took *C* = 2 so that syllable instances could take on speeds from “half as fast” to “twice as fast” as the base syllable.

#### Compute power

The models in section 5 (MoSeq dataset) were trained on a CPU cluster. As a comparison, a T-WARHMM with 30 discrete states and 31 warping variables took approximately 2.5 hr to train. A GP-WARHMM with the same parameters took approximately 15 hr to train. An ARHMM with 30 discrete states took approximately 1.5 hr to train.

## C Societal impacts

Research on obtaining quantitative characterizations of natural behavior will have broad implications for basic research in the field of neuroscience and neuroethology. Beyond basic neuroscience research, such methods can also be used to better characterize disease phenotypes and therefore also represent an important research direction for clinical sciences. One potential negative societal impact could come if these behavioral analysis approaches are applied to human surveillance, especially as they relate to identifying the influence of drugs on behavior.

